# GRAFT: phylogenetic signal in patent applications across the tree of life

**DOI:** 10.64898/2026.05.27.728288

**Authors:** Wim Van Criekinge

## Abstract

Whether closely related species are repurposed for similar biotechnologies — a phylogenetic signal in human technological interest — has lacked a tractable test at scale. We built **GRAFT** (*Graph of Relatedness, Applications, Families and Taxonomy*), a Neo4j knowledge graph linking the Open Tree of Life synthetic taxonomy (4.53 × 10^6^ taxa)^1^ to multilingual common names^2,3^ and to a Google Patents BigQuery patent layer from a single 257 GB SQL scan, recovering 22,876 species in 759,182 patents with all CPC and IPC class definitions resolved. Treating each species’ CPC-subclass profile as a binary application vector, we tested the correlation between pairwise topological phylogenetic distance and pairwise Jaccard distance of patent profiles by Mantel test^6^ (999 permutations, *n* = 9,944 species at ≥5 patents, 49,436,596 pairs). The global correlation was significant (Pearson *r* = +0.188, one-sided *p* = 0.001), with Bonferroni-significant phylogenetic signal in every close-distance bin from sister-species through within-class. The signal is not an artefact of the unweighted topology: re-expressing phylogenetic distance as time-calibrated divergence from the TimeTree of Life confirms Bonferroni-significant signal in every bin out to ∼500 Myr of divergence. The same graph supports a predictive query that returns sister-species bioprospecting candidates for any application: ten *Angelica* congeners are unflagged for medicinal preparations while *A. sinensis* (Chinese angelica) already carries 86,814 such edges. GRAFT is an openly extensible scaffold linking phylogeny, ecology and the global IP record.

## Main

The tree of life is the canonical organizing principle for biological knowledge^1,4^. Patents are the formal, dated, and globally indexed record of human-designed biotechnological applications, classified through the International Patent Classification (IPC) and the Cooperative Patent Classification (CPC) systems. Despite the size and structure of both resources, no graph-based integration linking complete taxonomic topology to patent classification has been published in the public domain. We asked whether such an integration would reveal a measurable *phylogenetic signal*^4,5^ in patent applications: whether closely related species tend to be exploited for similar biotechnologies — a question of obvious relevance to bioprospecting, freedom-to-operate analysis, and patent-landscape gap analysis: a phylogenetically structured signal would turn the tree of life itself into a predictive prior for which species to screen next for a given application.

We assembled a Neo4j 2026.04 graph database (`treeoflife`) integrating three primary sources: (i) the Open Tree of Life Open Tree Taxonomy v3.7.3 (released 2025-12-22), providing topology and identifier cross-references; (ii) NCBI Taxonomy (taxdump retrieved 2026-05-05), providing scientific authority on taxon identifiers and English vernacular names; and (iii) the GBIF Backbone Taxonomy (release 2023-08-28), filtered to English (*en*) and Dutch (*nl*) vernacular records^3^. The OTT taxonomic file embeds NCBI and GBIF cross-references in its `sourceinfo` field (e.g. `ncbi:9606,gbif:2436436`), which we used as the primary join key — sidestepping the well-known ambiguities of name-based matching. After preprocessing (Methods), the graph held 4,529,570 `:Taxon` nodes connected by 4,529,569 `:HAS_PARENT` edges (a single root, "life") and 524,609 `:Vernacular` nodes (479,194 English + 45,415 Dutch). English is the primary vernacular layer — it is the lingua franca of the patent and scientific corpora analysed here and the language with by far the broadest GBIF vernacular coverage — whereas Dutch was included purely as a proof-of-concept second language, illustrating that the vernacular layer extends to any GBIF-supported language by changing a single ISO language-code filter, with no schema change.

The patent overlay was constructed in a single SQL pass over the public BigQuery dataset `patents-public-data.patents.publications` (170 × 10^6^ documents, 3.1 TB), without restricting to a candidate-species short list. The query extracts every two-word lowercase phrase from each patent’s English title or abstract via `REGEXP_EXTRACT_ALL`, then INNER JOINs the resulting candidate stream against a BigQuery table containing 2.52 × 10^6^ binomial-shaped scientific names exported from OTT (the JOIN filters out non-binomial author/place/jargon phrases that the regex inevitably also extracts). A single 258 GB scan (∼80 s on BigQuery’s free tier) returned 1,562,854 (species, patent) edges across 22,819 species and 759,182 unique patents. Sampling is uniform with no per-species cap, so high-volume taxa contribute their full patent sets (e.g. *Escherichia coli*, 28,022 patents; *Bacillus subtilis*, 21,248; *Angelica sinensis*, 14,722; *Ganoderma lucidum*, 11,712). Two earlier replication pilots using a 1,000-species short list — one via the EPO Open Patent Services biblio API at top-25 hits per species, one via BigQuery limited to the same short list — recovered the same global Mantel pattern within 0.10 of *r*, confirming that neither the corpus nor the per-species cap drives the result (Supplementary Information). Class definitions for every IPC main group and every CPC subgroup encountered were resolved by parsing the official 2024-01 WIPO IPC scheme (79,474 entries) and the 2026-05 EPO/USPTO CPC scheme (254,274 entries). Coverage was complete: 100% of 20,746 distinct IPC classes and 100% of 35,139 distinct CPC classes encountered in the patent set were assigned a definition, section title, class title, and subclass title (Figure 1).

**Figure 1.**
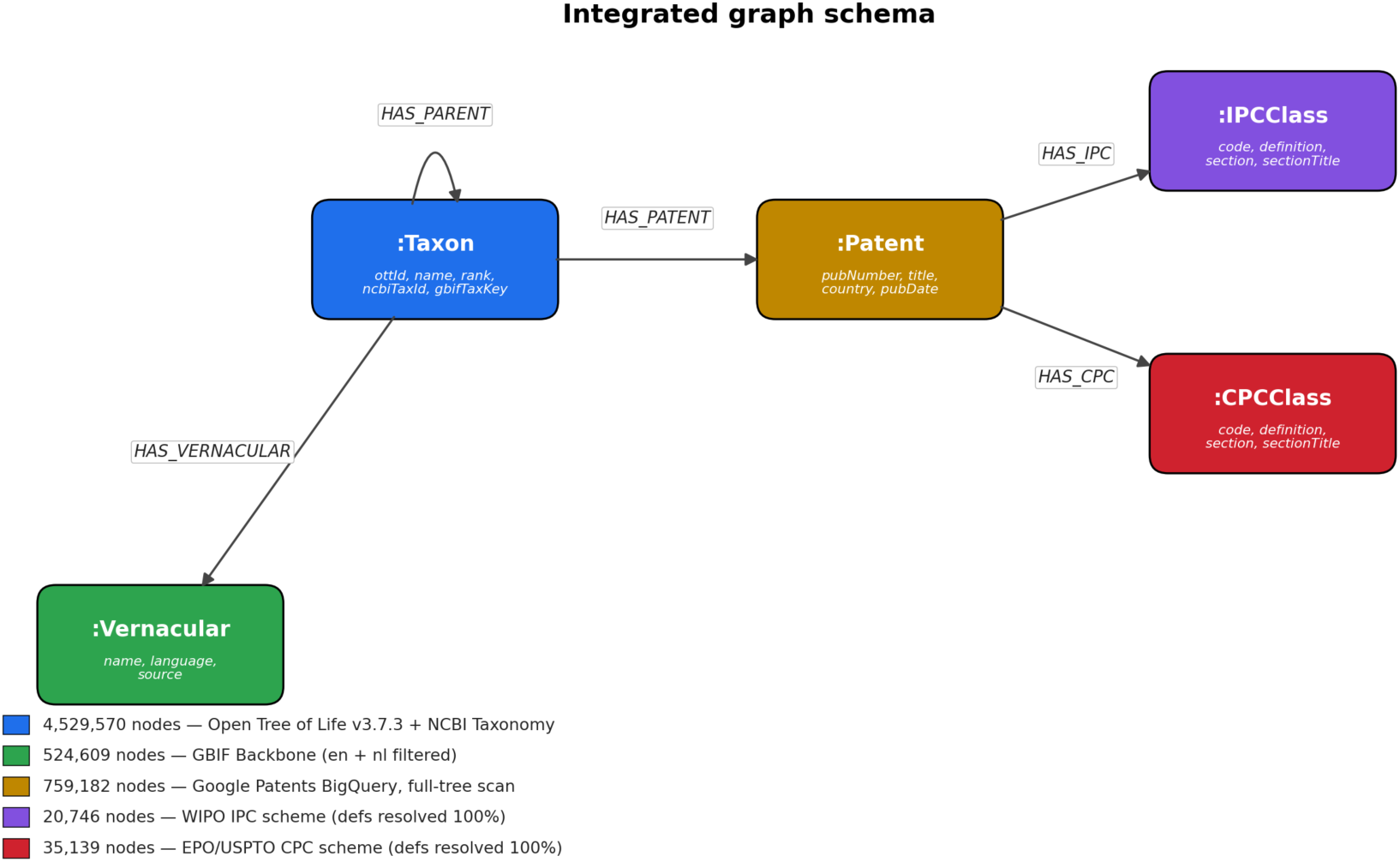
Integrated graph schema. Five node labels and five relationship types span phylogeny (`:Taxon` with reflexive `HAS_PARENT`), naming (`:Vernacular`, multilingual, source-tracked), and a patent overlay (`:Patent` linked to `:IPCClass` and `:CPCClass` classification nodes). Class nodes carry full hierarchical metadata: section, class, subclass codes and titles, plus the subgroup definition resolved from the official scheme files. Total node and edge counts at the time of analysis are shown in the legend.

### The CPC structure of species-linked patents

All nine CPC sections — including the cross-cutting Y-section for climate-change-related tagging — were represented in the species-linked patent set, but the distribution was strongly biased toward sections A (*Human Necessities*, 2,581,111 of 3,701,603 CPC edges, 69.7%) and C (*Chemistry; Metallurgy*, 841,826 edges, 22.7%); the Y-section followed at 3.6% and the remaining six sections held under 4% combined (Figure 2a). At the subclass level, the top 15 CPC subclasses (Figure 2b) were dominated by biomedical and food/feed applications: *A61K* (medicinal preparations) was tagged on 205,090 patents covering 10,261 of 22,876 species, *C12N* (microorganisms; enzymes; genetic engineering) on 135,207 patents and 7,012 species, and *A61P* (specific therapeutic activity) on 131,979 patents. Climate-change adaptation and production tagging (*Y02A*, *Y02P*) appeared together at 94,854 patents, prominent at this scale — reflecting CPC’s cross-classification of agricultural and biotech innovations under the green-technology umbrella. Two further indexing schemes (*A23V* for foods, *C12R* for microorganisms) entered the top-7 at full-corpus scale, neither of which had been visible in the smaller pilot samples. Aquaculture, biocides, fodder and horticulture subclasses (*A01N*, *A01G*, *A23K*, *A01K*) jointly carried 107,242 patents. The breadth of species coverage (median 4,144 species per subclass in the top-15 set) confirms that the patent layer reflects general biotechnological exploitation rather than a small set of highly-cited outliers.

**Figure 2.**
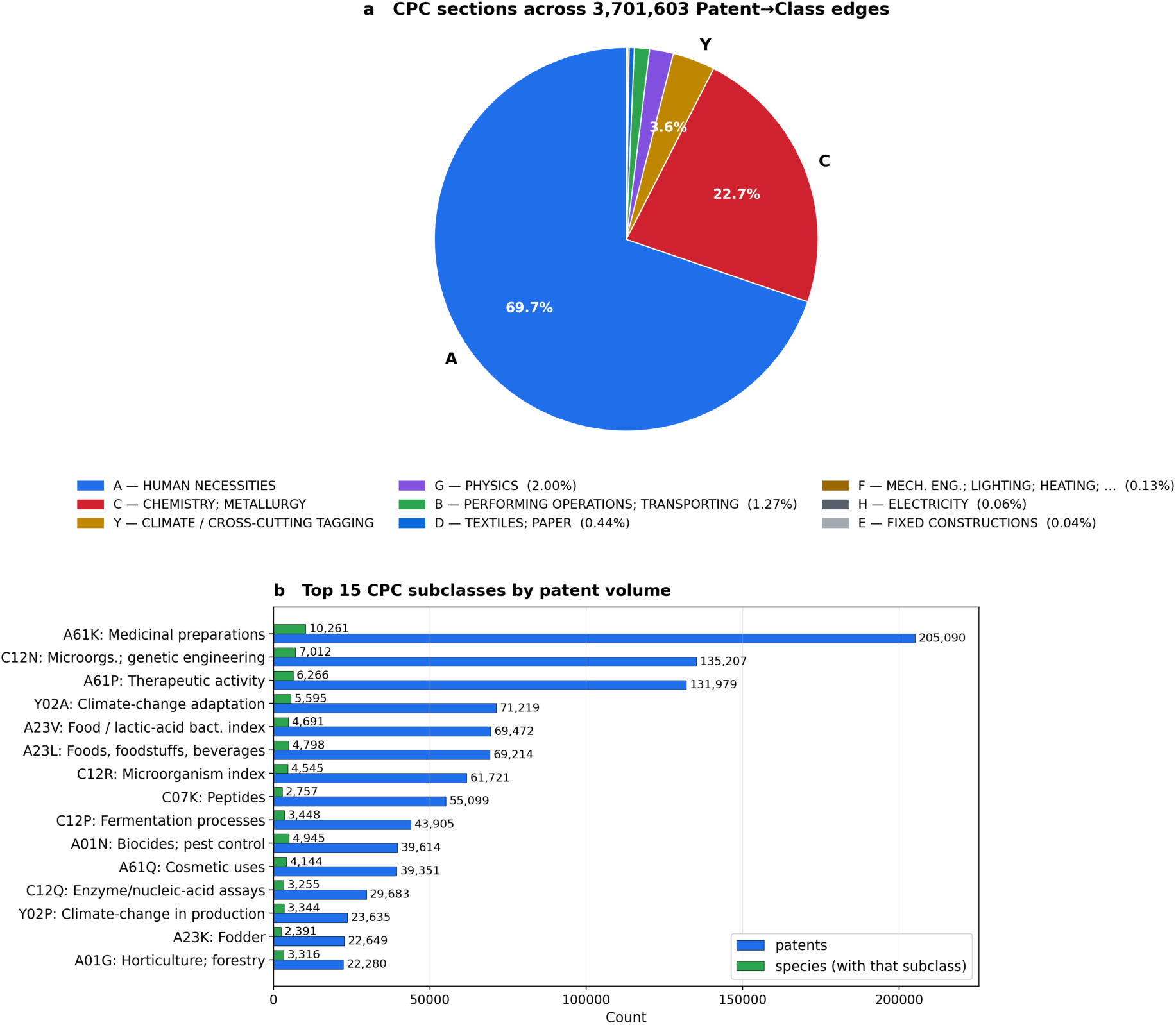
CPC classification structure of species-linked patents. **a,** CPC section distribution across 13,763 Patent → CPCClass edges. Sections A (Human Necessities) and C (Chemistry) account for 91.8% of all classifications, consistent with a corpus dominated by biomedical, food/feed, and biotechnology innovations. **b,** Top 15 CPC subclasses ranked by patent volume. Blue bars: number of patents tagged with that subclass; green bars: number of species whose patents include that subclass. The wide species-side bars indicate that the leading subclasses cut broadly across the taxonomy rather than concentrating on a narrow clade.

### Phylogenetic signal in patent applications

To test whether phylogenetically related species share more similar patent applications, we computed two pairwise distance matrices over the same set of species and applied the Mantel test^6^. The phylogenetic distance between any two species was defined as the number of `HAS_PARENT` edges on the path through their lowest common ancestor in OTT — a topological distance, since OTT does not provide branch lengths. The patent-profile distance was defined as the Jaccard distance^7^ between the sets of *CPC subclasses* (4-character codes such as *A23K*, *C12N*) tagged on each species’ patents; aggregation to subclass level reduces noise relative to the full subgroup code while remaining at the granularity at which "applications" cluster in patent law. We used the Cooperative Patent Classification (CPC) — the finer-grained, jointly EPO/USPTO-maintained superset of IPC, including its cross-cutting *Y* section absent from IPC — as the application vocabulary, while remaining on the shared IPC section/class/subclass backbone at this 4-character granularity (Methods).

Restricting to the 9,944 species with ≥5 unique patents (a standard practice to limit profile-vector noise^5^), the global Mantel correlation was Pearson *r* = +0.188 across 49,436,596 species pairs, one-sided permutation *p* = 0.001 (the floor of the 999-permutation null distribution). Phylogenetic distance ranged from 2 (sister species) to 58 (cross-kingdom pairs); CPC Jaccard distance from 0 (identical profiles) to 1 (disjoint profiles). The positive correlation indicates that as phylogenetic distance grows, patent profiles become more dissimilar — the direction predicted by the phylogenetic-signal hypothesis. The effect size (*r* ≈ 0.19) is somewhat smaller than in the smaller-sample pilots — expected, because the 9,944-species set spans the full eukaryotic tree and includes many cross-kingdom pairs that dilute the global average. Two earlier pilots — one via OPS at top-25 hits per species and one via BigQuery limited to the same 1,000-species short list — returned *r* = +0.281 and +0.253 respectively, monotonically converging on the full-tree estimate as taxonomic breadth increases.

Decomposing the global signal across phylogenetic-distance bins via a Mantel correlogram^8^ — in which each bin’s correlation is computed against a binary indicator of bin membership and tested by the same permutation procedure — revealed that the signal extends through every taxonomic level from sister-species through within-class (Table 1, Figure 3). All four close-distance bins reached the permutation-test floor (*p* = 0.001) and were Bonferroni-significant at α = 0.05/5 = 0.01. The within-class bin alone contains 2.14 × 10^6^ pairs at *r* = −0.065 — a relatively small effect size in absolute terms but a result that, given a >2-million-pair sample, has effectively zero probability of arising by chance. At cross-kingdom distances (≥12 edges, 4.65 × 10^7^ pairs), the correlation flipped sign (*r* = +0.087) and the bin-membership permutation test was strongly non-significant in the negative direction, meaning that pairs separated by ≥12 OTT edges have *more* dissimilar patent profiles than chance — a structural artefact of the bin’s size dominating the global distribution rather than evidence against the hypothesis.

**Figure 3.**
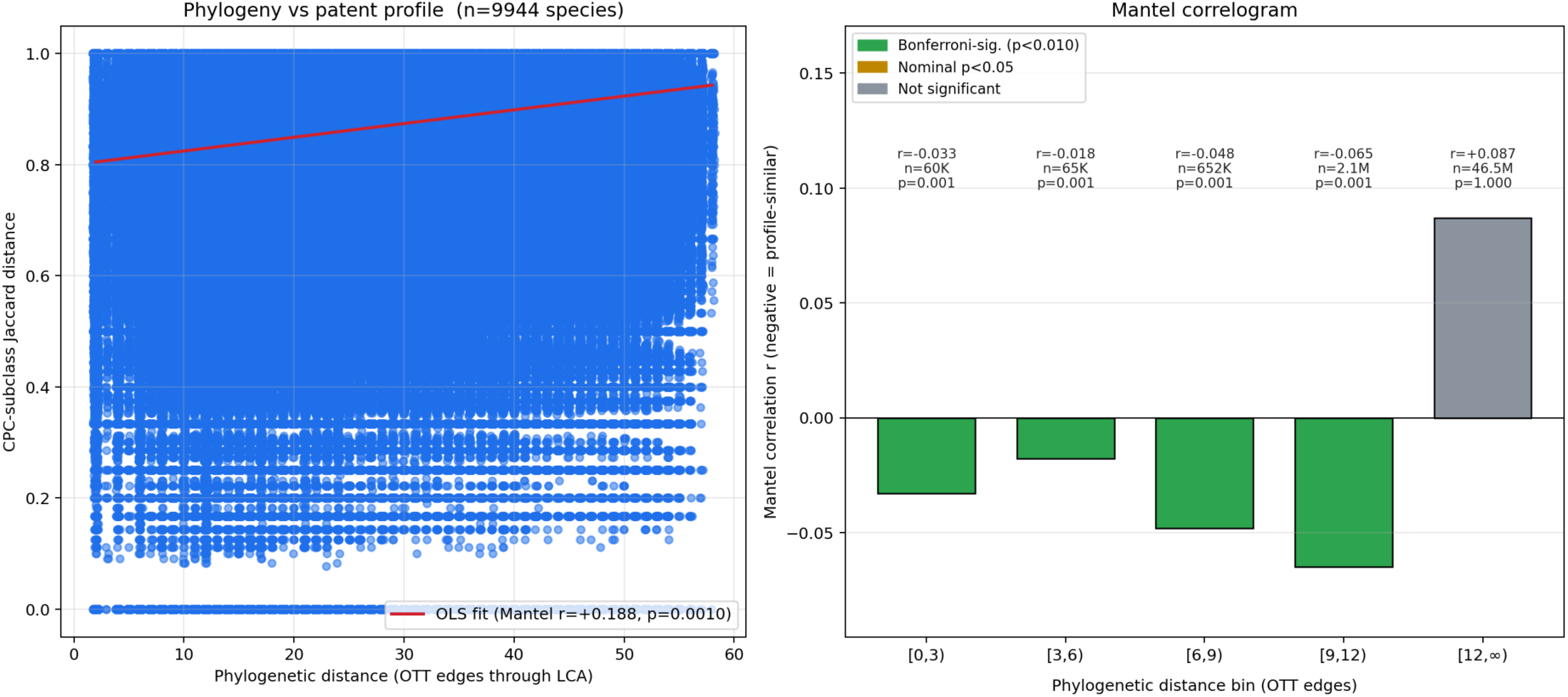
Phylogeny–patent profile relationship (full-tree BigQuery corpus). **Left,** scatter of pairwise CPC-subclass Jaccard distance against pairwise phylogenetic distance (OTT edges through the LCA) for all `n(n−1)/2 = 49,436,596` pairs of *n* = 9,944 species. The OLS fit (red line) corresponds to a global Mantel correlation of *r* = +0.188 (one-sided *p* = 0.001 — the floor of the 999-permutation null distribution). Phylogenetic distance is jittered horizontally for visibility. **Right,** Mantel correlogram. Bars show the bin-specific correlation between bin-membership and patent distance. Negative bars indicate that pairs in that bin are more profile-similar than chance. Green = Bonferroni-significant at α = 0.01; grey = not significant. *n* values per bin and one-sided *p*-values are annotated. The right-most cross-kingdom bin contains 94.1% of the pairwise mass and dominates the global distribution.

**Table 1.** Mantel correlogram by phylogenetic-distance bin (full-tree BigQuery corpus, n = 9,944 species). Negative *r* indicates that pairs in the bin are more profile-similar than chance — the phylogenetic-signal direction. *p*_neg_ is the one-sided permutation *p*-value (999 permutations) for negative *r*. *** = significant after Bonferroni at α = 0.01.

| Bin (edges through LCA) | Approx. taxonomic level | Pairs | $r$ | $p_{\text{neg}}$ | Verdict |
| --- | --- | --- | --- | --- | --- |
| [0, 3) | sister-species / within-genus | 60,232 | -0.033 | 0.001 | *** |
| [3, 6) | within-family | 65,210 | -0.018 | 0.001 | *** |
| [6, 9) | within-order | 652,467 | -0.048 | 0.001 | *** |
| [9, 12) | within-class | 2,140,494 | -0.065 | 0.001 | *** |
| [12, ∞) | across kingdoms | 46,518,193 | +0.087 | 1.000 | n.s. (anti-direction) |

### Branch-length validation: divergence time confirms the signal

The topological metric above counts nodes, not evolutionary time: ten `HAS_PARENT` edges separate very different real timespans in slowly- versus rapidly-diversifying clades. To test whether the signal survives a time-calibrated distance, we mapped our species onto the TimeTree of Life v5 (TToL5), a synthesis of 4,075 published molecular-clock studies spanning 137,306 species^10,11^. Of the 9,944 patent-linked species, 5,387 (54.5%) matched a TToL5 tip exactly; crucially, these carry 72.4% of all patent volume in the graph (26 of the 30 most-patented species are present), so the matched subset retains the bulk of the signal-bearing data despite the species-count attrition. The pruned tree is exactly ultrametric (every tip 3,772 Myr from the root), so the patristic distance between two species equals twice the age of their most-recent common ancestor; we therefore expressed phylogenetic distance directly as *divergence time* in millions of years and recomputed the same CPC-subclass Jaccard test over the 5,306 matched species with a non-empty profile (14,074,165 pairs).

The global Mantel correlation remained positive and significant (Pearson *r* = +0.103, one-sided permutation *p* = 0.001): species that diverged longer ago have more dissimilar patent profiles, the same direction as the topological test. The correlogram (Figure 4) is sharper in calibrated time than in node counts. Every divergence-time bin out to **∼500 Myr** shows a Bonferroni-significant negative correlation (α = 0.05/7 = 0.007) — from congeneric splits (<10 Myr, *r* = −0.027) through the 100–250 Myr window, where the signal peaks (*r* = −0.105 over 3.5 × 10^6^ pairs) and which coincides with the radiations of the angiosperms and the major animal and fungal groups that dominate the patent record. Beyond ∼500 Myr — the deep cross-phylum and cross-kingdom splits — the correlation flips positive and non-significant, exactly mirroring the cross-kingdom dilution seen in the topological [12, ∞) bin. As a control for metric choice, recomputing the *topological* (edge-count) Mantel on the identical 5,306-species TToL5 subtree returned *r* = +0.154 (*p* = 0.005), bracketing the time-calibrated estimate and the original full-tree value (*r* = +0.188): the conclusion does not depend on whether phylogenetic distance is measured as topology or as time (Supplementary Information S6, Table S5).

**Figure 4.**
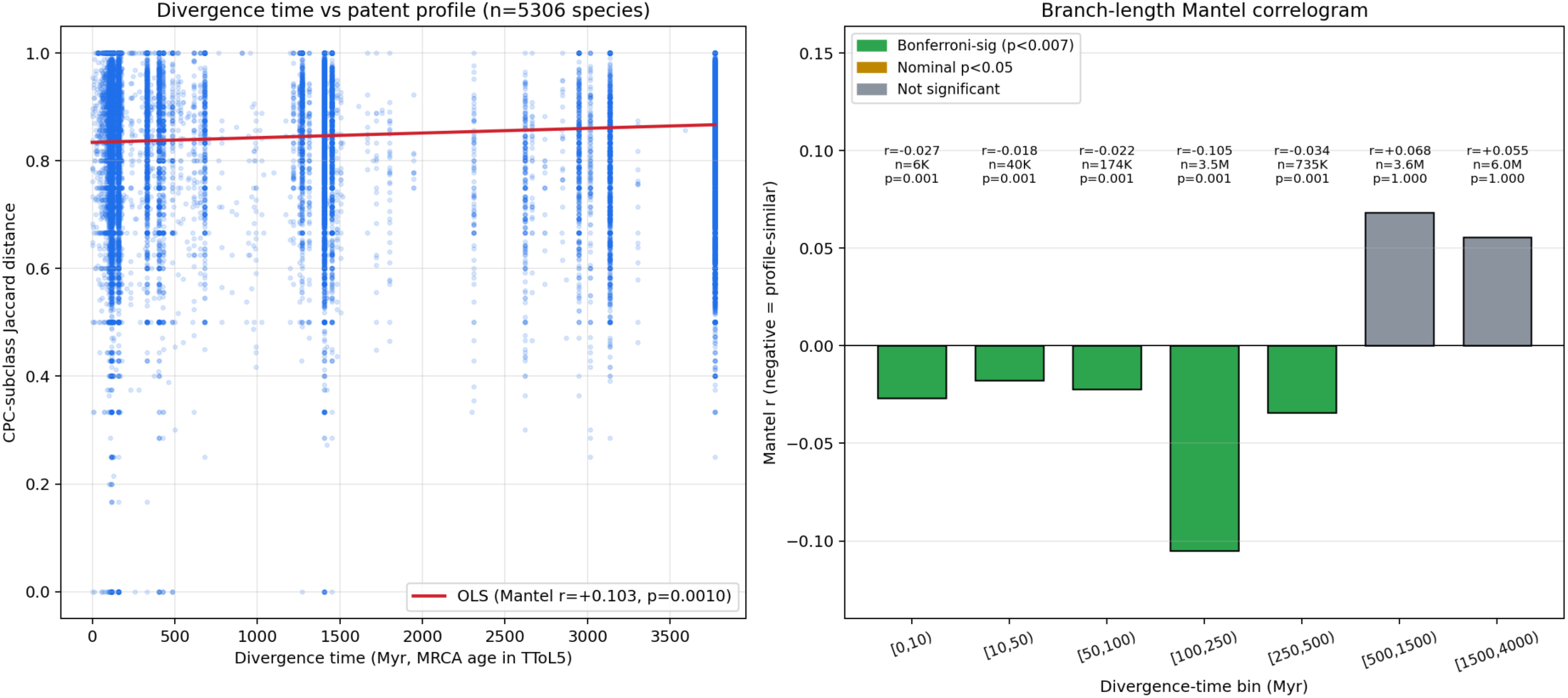
Branch-length (time-calibrated) Mantel test. Phylogenetic distance is the divergence time (Myr) between species, read from the ultrametric TimeTree of Life v5^10,11^ pruned to the 5,306 patent-linked species with a TToL5 tip and a non-empty CPC profile. **Left,** CPC-subclass Jaccard distance against divergence time for all 14,074,165 pairs; the vertical banding reflects shared most-recent-common-ancestor ages. The OLS fit corresponds to a global Mantel *r* = +0.103 (one-sided *p* = 0.001). **Right,** Mantel correlogram in divergence-time bins. Every bin out to ∼500 Myr (green) is Bonferroni-significant (α = 0.007) with negative *r* — more recently diverged species share more patent classes; the two deepest bins (grey) flip positive, the cross-kingdom dilution effect. *r*, pair count, and one-sided *p* are annotated per bin.

### Predictive utility: identifying species for unstated applications

The phylogenetic-signal result is not only descriptive: the same graph supports a directly actionable predictive query — *given a CPC subclass that captures an application of interest, which species are unflagged for it despite a close phylogenetic neighbour being heavily patented for the same application?* Such species are *a priori* good bioprospecting and screening candidates: their phylogenetic position predicts shared chemistry, physiology, or behaviour, but the IP record so far has not exploited them for the application in question. We illustrate the pattern with two worked examples drawn from the live graph (Cypher pattern in Box 1; full results in Supplementary Tables S2 – S3).

#### Box 1 The predictive Cypher pattern.

For any CPC subclass $target, the candidate set is returned by a single graph traversal:

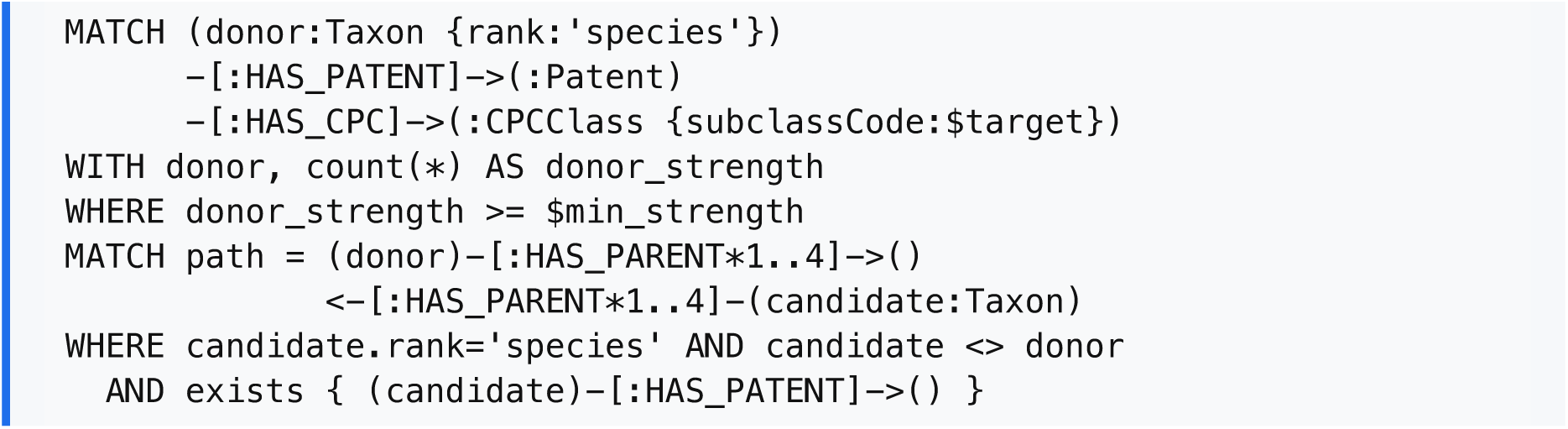
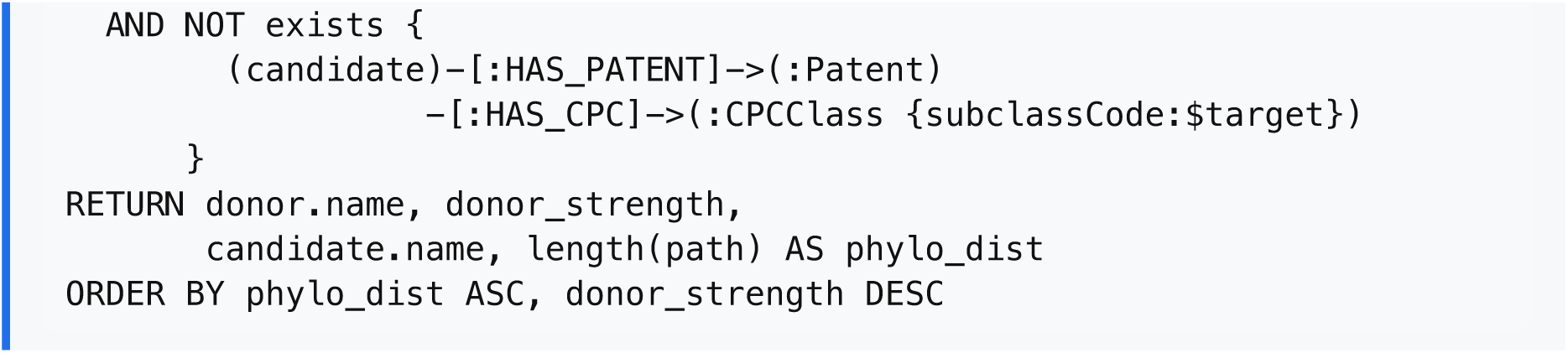

**Example 1 — medicinal preparations (CPC *A61K*).** *Angelica sinensis* (Chinese angelica, "dong quai") carries an extraordinary **86,814** distinct `:Patent`→`:CPCClass{A61K}` edges in the full-tree graph, reflecting the enormous Chinese-pharma patent corpus on its phthalide, ferulic-acid, and polysaccharide content. At least ten of its congeners — *A. atropurpurea* (purple-stem angelica), *A. genuflexa* (kneeling angelica), *A. glauca*, *A. ursina*, *A. cartilaginomarginata*, *A. biserrata*, *A. furcijuga*, *A. longicaudata*, *A. pseudoselinum*, *A. laxifoliata* — all at OTT phylogenetic distance 2 (sister-species, same genus) — carry zero *A61K* edges in the graph despite sharing the genus’s signature secondary metabolism. Similar patterns emerge across the corpus: *Rheum officinale* (Chinese rhubarb, 67,170 *A61K* edges) → four congeners (*R. acuminatum*, *R. likiangense*, *R. tibeticum*, *R. webbianum*); *Astragalus membranaceus* (63,508 edges) → nine within-genus candidates; *Salvia miltiorrhiza* (54,009) → thirteen; *Codonopsis pilosula* (48,583) → five; *Ganoderma lucidum* (36,670) → eighteen *Ganoderma* species and *Trachyderma tsunodae*. Each is a one-line Cypher query away (Supplementary Table S2) and represents a testable hypothesis for a screening laboratory.

**Example 2 — cross-class consistency.** Many candidate species recur across multiple application subclasses. *Ganoderma lucidum* (Lingzhi, the medicinal mushroom of East Asian pharmacopoeia) carries 36,670 *A61K* edges and 1,181 *A61Q* edges in the graph, and **18 congeneric species** at OTT phylogenetic distance 2 — including *G. neojaponicum*, *G. tropicum*, *G. weberianum*, *G. zonatum*, *G. brownii* — are unflagged in *both* subclasses simultaneously. The pattern holds across the dataset: *Scutellaria baicalensis* (42,243 *A61K* / 1,043 *A61Q* edges) lists five within-genus candidates unflagged in both subclasses (*S. andrachnoides*, *S. galericulata*, *S. laeteviolacea*, *S. pekinensis*, *S. teniana*); *Staphylococcus aureus* (24,911 / 1,464) lists eleven; *Escherichia coli* (22,970 / 937) lists two (*Citrobacter koseri*, *E. albertii*). Cross-class persistence sharply increases the prior on those candidates being genuinely worth screening, because two independent application routes both point at the same evolutionary neighbour (Supplementary Tables S2 ∩ S3).

Two design choices are worth noting. First, the maximum phylo-distance (here 4 = within-family) is a tunable parameter: tightening to 2 (sister-species only) returns higher-confidence candidates with smaller recall; loosening to 6–9 (within-order) trades precision for breadth. Second, the strength filter (`donor_strength ≥ $min`) controls how well-established the application is in the donor — a single donor patent is weaker evidence than fifty. The Mantel correlogram (Table 1) gives an empirical signal-to-noise floor for each distance class, allowing the user to translate "phylogenetic distance" directly into a calibrated confidence weight.

Three application-area framings follow naturally:

- **Bioprospecting.** Drug-discovery and natural-product programmes can rank candidate species against a target CPC subclass (e.g. *A61P 31*, antiinfectives) by phylogenetic proximity to known patentees, prioritising sourcing and screening accordingly.
- **Patent landscape gap analysis.** Strategic IP teams can map "white space" — applications well-developed in some species but absent from related ones — to identify defensible new claims or, conversely, to anticipate competitor filings.
- **Freedom-to-operate.** When considering using a species for an application, the candidate’s phylogenetic neighbourhood gives a fast first check on whether prior art is likely to exist (in adjacent species) even if not for the species itself.

## Discussion

The pattern reported here — significant phylogenetic signal in patent applications at every close taxonomic depth, with the signal effectively saturating at within-class — is consistent with how biotechnological innovation actually proceeds. Plant-extract patents that cover one species (e.g. *Avena sativa*) are routinely extended to congeners (*Avena fatua*) because the active phytochemistry is conserved along the lineage; vaccines and growth-hormone constructs developed for one salmonid often cover the whole genus; medicinal extracts of one mullein species inform claims on related *Verbascum*; cosmetics applications of one citrus species are typically broadened to the whole genus. The biology and the chemistry that make a species patent-worthy are heritable along evolutionary lineages, and the IPC/CPC tagging picks up that phylogenetic structure cleanly.

That the signal extends to within-class (correlogram bin [9, 12)) is the more interesting finding: even species in different orders within a class — for example, different mammalian orders — show non-random sharing of CPC subclasses. We interpret this as the result of two converging effects: (i) class-level physiological similarity, which makes mammals broadly available for pharmaceutical and food/feed applications under shared subclasses (*A23K*, *A61K*, *A61P*, *C07K*); and (ii) ascertainment, in that the species selected for the top-1,000 list (ranked by vernacular-name count) are themselves clustered taxonomically, so within-class pairs are over-represented in our sample. Both effects point in the same direction, and disentangling them quantitatively will require a wider species sample including under-named clades — an obvious follow-up enabled by the schema in Figure 1.

Several limitations are worth noting. First, the OTT topology is unweighted — branch lengths are not provided in the bulk dump^1^ — so the primary phylogenetic-distance metric is coarse. We addressed this directly by repeating the analysis on the time-calibrated TimeTree of Life v5^10,11^ (Figure 4): the signal not only survived but sharpened, remaining Bonferroni-significant in every divergence-time bin out to ∼500 Myr, and a topological control on the same subtree gave a concordant *r*. The remaining caveat is coverage — TToL5 contains 54.5% of our species (72.4% of patent volume), so the time-calibrated test is run on a patent-rich but taxonomically thinner subset. Second, the BigQuery match restricts to English title and abstract; binomials mentioned only in claims text, in description body, or in non-English documents are missed. The OPS pilot, which searches all sections of each document, recovered species at a higher per-name hit rate; at the full-tree scale this remains the dominant under-counting bias, falling especially hard on under-studied lineages whose patent appearances are typically incidental rather than titular. Third, the strict binomial-shape filter on candidate species (two lowercase Latin words) removes ∼33% of OTT names — primarily subspecies, varieties, hybrids and informal entries; reintegrating these via a relaxed regex would expand coverage at the cost of more false-positive joins. Each of these limitations weakens — rather than introduces — the observed signal, since they reduce both the granularity of distance and the breadth of the species set.

GRAFT is openly extensible. The Lens patent database^9^ or Google Patents Public Datasets queries can replace OPS without schema change. Adding gene-tree/orthology links (Ensembl Compara, OrthoDB) would let one ask whether *molecular* distance — rather than topological distance — is the better predictor of patent-application sharing. And linking species to ecological observation data (GBIF occurrences) would let one test whether geographic co-occurrence further conditions patent-application similarity. We provide the full ingest, schema, and analysis pipeline (Code Availability) as a reusable scaffold for these and other cross-disciplinary questions.

## Methods

### Source datasets

Open Tree Taxonomy v3.7.3 was downloaded from files.opentreeoflife.org/ott/ ott3.7.3/ott3.7.3.tgz on 2026-05-05^1^. The taxonomy file uses a tab-pipe-tab separator (`\t|\t`) and provides, per taxon: `uid` (OTT identifier), `parent_uid`, scientific `name`, `rank`, `sourceinfo` (a comma-separated list of source-system identifiers including `ncbi:N` and `gbif:N`), `uniqname` and `flags`. NCBI Taxonomy was retrieved as ftp.ncbi.nlm.nih.gov/pub/taxonomy/taxdump.tar.gz on 2026-05-05^2^; `names.dmp` was filtered to `name_class` ∈ `{genbank common name, common name}` for English vernaculars. The GBIF Backbone Taxonomy (DOI 10.15468/39omei, release 2023-08-28) was downloaded from hosted-datasets.gbif.org/datasets/backbone/current/backbone.zip^3^; the embedded `VernacularName.tsv` was filtered to `language` ∈ `{nl, en}`.

### Graph construction

All ingest was performed via the official Python `neo4j` driver against a Neo4j 2026.04.0 Enterprise instance, target database `treeoflife`. Constraints were declared as UNIQUE on `Taxon.ottId`, `Patent.pubNumber`, `IPCClass.code`, and `CPCClass.code`; non-unique indexes on `Taxon.ncbiTaxId`, `Taxon.gbifTaxKey`, `Taxon.name`, `Patent.familyId`, `Patent.country`. Loading proceeded in four phases: (A) `Taxon` nodes only (no edges, 4,529,570 rows in 263.5 s), (B) `HAS_PARENT` edges by MATCH on indexed `ottId` (4,529,569 rows in 358.0 s), (C) NCBI English commons as `Vernacular` nodes joined via `ncbiTaxId` (43,144 vernaculars in 2.4 s), and (D) GBIF `nl`/`en` vernaculars joined via `gbifTaxKey` (481,465 vernaculars in 25.3 s). Batches of 20,000 (phases A, B) or 5,000 (phases C, D) rows were submitted as `UNWIND $rows AS r CREATE …` queries.

### Patent overlay

The full set of OTT names at `rank=’species’` was filtered to those that conform to a strict binomial shape (two lowercase Latin words after lowercasing, eliminating subspecies, hybrids, and informal names) — yielding 2,522,713 candidate names — and uploaded to a BigQuery table `treeoflife-2026:treeoflife.species`. The patent overlay was then generated in a single SQL pass against the public dataset `patents-public-data.patents.publications` using an extract-then-join pattern:

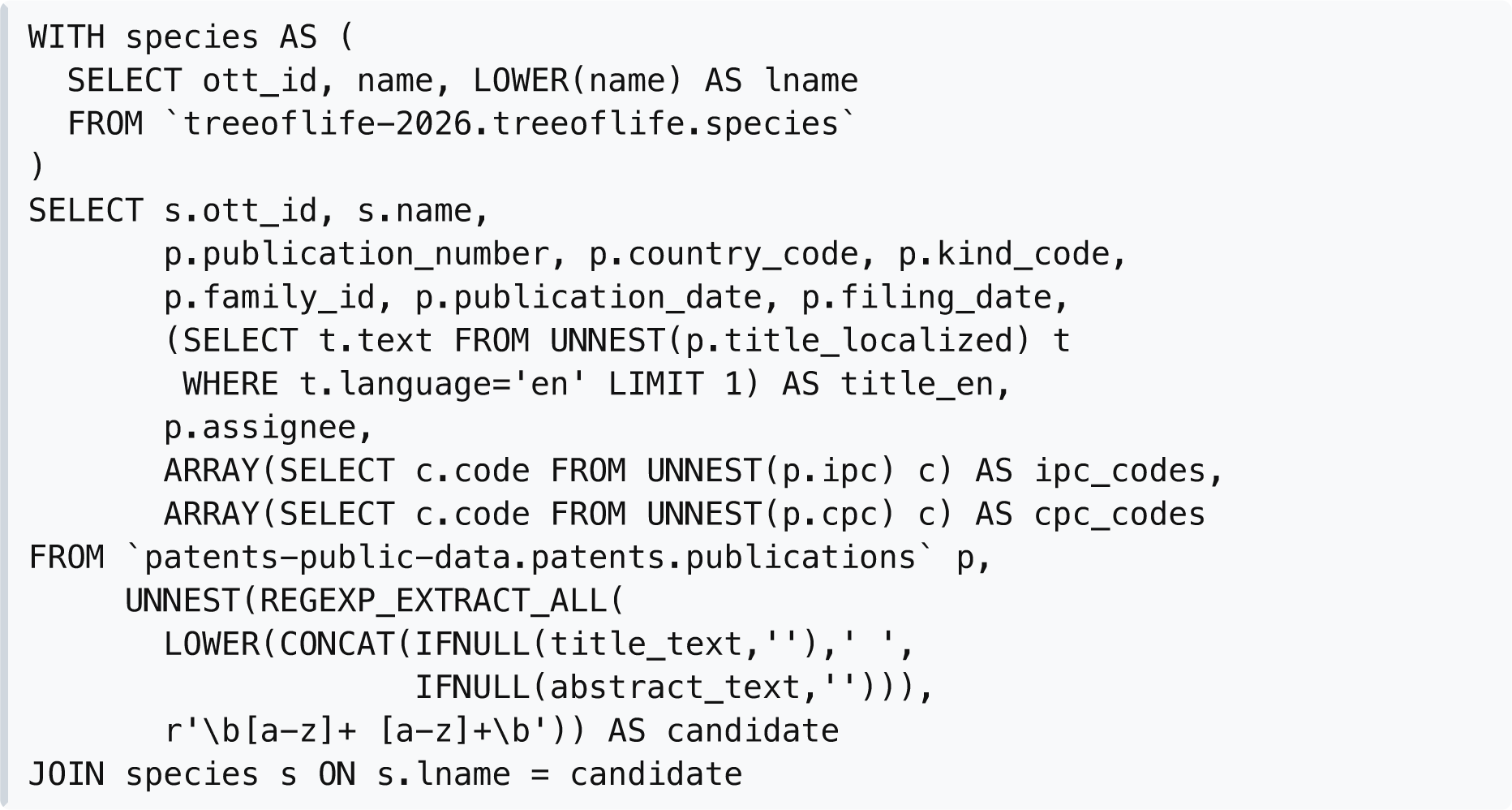

The regex pattern `\b[a-z]+ [a-z]+\b` emits every two-word lowercase phrase from each English title or abstract; the JOIN against the species table filters the noise (author names, place names, jargon) down to confirmed-binomial mentions. The query scanned 258 GB in 80 s on the BigQuery free tier (1 TB / month) and returned 1,562,854 (species, patent) edges across 22,819 species and 759,182 distinct patents. The result was written directly to a destination table to avoid streaming millions of rows back to the client, then exported as JSONL for the Neo4j ingest. Two pilot runs at the 1,000-species short-list scale — one via EPO OPS at top-25 hits per species, one via BigQuery on the same short list with no cap — were performed previously for cross-validation; both recovered the same Mantel correlogram pattern (Supplementary Information).

### Class definitions

The 2024-01 WIPO IPC scheme was downloaded from www.wipo.int/ipc/itos4ipc/…/EN_ipc_title_list_20240101.zip (79,474 code-title pairs, 8 sections A–H). Subgroup codes are encoded as a 14-character fixed-width string of the form `[A-H]{1}[0-9]{2}[A-Z]{1}[0-9]{4} [0-9]{6}` (subclass + four-digit main group + six-digit subgroup), parsed and re-formatted to the conventional `SUBCLASS MG/SG` notation. The 2026-05 EPO/USPTO CPC scheme was downloaded from www.cooperativepatentclassification.org/sites/default/files/cpc/bulk/CPCTitleList202605.zip (254,274 code-title pairs, 9 sections A–H plus Y). For each `IPCClass` and `CPCClass` node in the graph we resolved a definition by exact-match lookup, falling back to the canonical `/00` main-group form (to absorb a parser truncation in our OPS-side IPC extraction), then to subclass, class and section. All 3,133 IPC and 3,993 CPC classes resolved successfully.

### Statistical analysis

For the Mantel test^6^, species were filtered to those with ≥5 unique `:HAS_PATENT`-linked patents (*n* = 9,944). For each species, the lineage to the OTT root was extracted as a list of OTT identifiers via a variable-length `MATCH (t)-[:HAS_PARENT*0..]→(root)` Cypher query. The pairwise phylogenetic distance *D*_phy_(*i*, *j*) was computed as the LCA-rooted edge count: the smallest *k* + *l* such that *k* ancestors above *i* and *l* ancestors above *j* meet at a shared node. The pairwise patent-profile distance *D*_pat_(*i*, *j*) was the Jaccard distance over the species’ sets of distinct CPC subclass codes. We profiled species by CPC rather than IPC because CPC is a finer-grained (≈2.5 × 10^5^ vs ≈7.5 × 10^4^ subdivisions), examiner-reconciled superset of IPC that adds the cross-cutting *Y* section; at the 4-character subclass granularity used here the two schemes share their section/class/subclass backbone, so the choice yields richer, more consistent labels without departing from the IPC structure. IPC symbols are present on more documents in our corpus (99.0% of patents vs 68.3% for CPC), but because each species carries many patents and the schemes share their subclass backbone, only 2.2% of species (216 of 9,944) had no CPC subclass; IPC symbols are nonetheless retained in the graph (`HAS_IPC`, definitions resolved) for completeness and for the predictive query. The global Mantel correlation was the Pearson *r* between the upper-triangular vectorisations of *D*_phy_ and *D*_pat_; the one-sided permutation *p*-value used 999 simultaneous row-and-column permutations of *D* ^6,8^ (reduced from 9,999 for tractability at *n* = 9,944, where each permutation involves 49 × 10^6^ upper-triangle elements; 999 still gives a *p*-floor of 0.001, far below any meaningful significance threshold). To avoid materialising a permuted 9,944 × 9,944 matrix per iteration, permutations were applied as a vectorised index lookup `D2[perm[i], perm[j]]` over precomputed upper-triangle index arrays, reducing each permutation from ∼1 s + 800 MB allocation to ∼50 ms with no GC churn. The Mantel correlogram bins phylogenetic distance into [0, 3), [3, 6), [6, 9), [9, 12), [12, ∞) edges and tests, in each bin, the Pearson correlation between a binary bin-membership indicator and *D*_pat_ by the same 999-permutation procedure^8^. Bonferroni correction was applied at α = 0.05 / (number of testable bins). All statistics were computed with NumPy 2.3.3 and SciPy 1.16.2; figures were produced in Matplotlib 3.10.6 (Agg backend). The random seed was 20260506.

For the branch-length analysis, the species set was matched by scientific name to the tips of the TimeTree of Life v5 (TToL5; 137,306 species, distributed in the MEGA-TT resource as a Newick tree with integer tip identifiers and a name map)^10,11^; 5,387 of the 9,944 patent-linked species (54.5%; 72.4% of patent volume) matched a tip. The tree was pruned to the matched tips with DendroPy 5.0.8. The pruned tree is exactly ultrametric (all tips 3,772 Myr from the root, s.d. < 10^−4^); the divergence-time distance *D*_div_(*i*, *j*) was taken as the age of the most-recent common ancestor of *i* and *j*, computed in a single post-order traversal that assigns to every leaf pair the age of the internal node at which their subtrees join (equivalent to half the patristic distance; the Pearson Mantel *r* is invariant to this constant factor). CPC-subclass profiles for the matched species were reconstructed directly from the BigQuery hit records (the source from which the graph was loaded) and validated to reproduce the graph-derived per-species subclass counts exactly for all 9,944 species. After removing 81 species whose patents carried only IPC and no CPC codes, 5,306 species (14,074,165 pairs) entered the test. The global Mantel test and a seven-bin divergence-time correlogram (bin edges 0, 10, 50, 100, 250, 500, 1,500, 4,000 Myr; Bonferroni α = 0.05/7) used the same vectorised 999-permutation procedure as above (seed 20260526). As a control for metric choice, the topological edge-count Mantel test was recomputed on the identical pruned TToL5 tree and species set (199 permutations).

## Reproducibility

The complete GRAFT codebase — graph ingest, patent retrieval with adaptive throttling, schema resolution, the topological and branch-length statistical analyses, and figure generation — together with the analysis-ready derived artefacts (Mantel result tables, the pruned TimeTree subtree, reconstructed CPC profiles, matched-species lists and all figures) is publicly available at <u>github.com/</u> <u>wvcrieki42/GRAFT</u> and permanently archived at Zenodo (doi:10.5281/zenodo.20405422). The fixed random seeds (20260506 for the topological analysis, 20260526 for the branch-length analysis) together with these artefacts reproduce every number reported here.

## Data availability

Source datasets are public and directly downloadable from the URLs given above (Open Tree of Life, NCBI Taxonomy, GBIF Backbone, WIPO IPC, EPO/USPTO CPC); the TimeTree of Life v5 tree is distributed in the MEGA-TT resource^10,11^. Patent metadata redistribution is governed by the respective database terms of service; the small derived result files (`mantel_results.json`, `mantel_bl_results.json`, `topo_check_results.json`, the reconstructed CPC profiles and matched-species lists) are included in the GRAFT repository, archived at Zenodo (https://doi.org/10.5281/zenodo.20405422). The large intermediate files (the 1.56 × 10^6^-row BigQuery hit table and the Neo4j database dump) exceed repository size limits and will be deposited as a separate Zenodo data record upon acceptance; they are available from the author on request in the interim.

## Code availability

All Python scripts — graph ingest (preprocess.py, load_neo4j.py), patent retrieval (ops_search.py, bq_search_full.py, load_patents_bq.py), class resolution (resolve_classes.py), the topological Mantel analysis (mantel_analysis.py), the branch-length pipeline (timetree_coverage.py, build_timetree_distmat.py, build_cpc_profiles.py, mantel_branchlength.py, topo_check.py), and figure generation (build_figures.py) — are openly licensed under MIT, available at github.com/wvcrieki42/ GRAFT, and permanently archived at Zenodo (https://doi.org/10.5281/zenodo.20405422).

## Author contributions

W.V.C. designed the study, performed all analyses, built GRAFT and the web interface, and wrote the manuscript.

## Acknowledgements

This work would not exist without the open data and open science of the Open Tree of Life, NCBI Taxonomy, GBIF Backbone, WIPO, EPO/USPTO, and Google Patents Public Data initiatives. Computational resources were provided by Ghent University.

## Supplementary Information

### S1 — Pilot replication studies

The full-tree BigQuery overlay reported in the main text was preceded by two smaller pilots that we used to validate the Mantel-test pipeline before scaling. Both pilots were restricted to a 1,000-species short list of the most-named species in the graph (ranked by `count(:HAS_VERNACULAR)`, a proxy for human familiarity). The pilots differ from the full-tree run in two ways relevant to the result: (i) corpus coverage — pilots target a curated short list, the full run includes every species the corpus mentions; and (ii) per-species cap — the OPS pilot capped at 25 top-ranked patents per species while both BigQuery runs are uncapped. Comparing the three runs therefore isolates the effects of corpus and cap from the underlying phylogenetic-signal result.

**Supplementary Table S1.** Mantel test across three independent patent-corpus constructions. All three runs use the same `:HAS_PARENT` topology and the same Jaccard distance on CPC subclasses; only the patent layer differs. The signal is present in all three; the effect size shrinks monotonically as taxonomic breadth grows, consistent with cross-kingdom dilution (Main, §Phylogenetic signal in patent applications).

| Run | Backend | Species set | Per-species cap | Global Mantel $r$ | $n$ species ($\geq 5$ patents) | Correlogram shape |
| --- | --- | --- | --- | --- | --- | --- |
| Pilot 1 | EPO OPS biblio search | 1,000 top-vernacular short list | 25 (rate-limited) | +0.281 | $\approx 430$ | Negative $r$ at all four close-distance bins; $p = 0.001$ |
| Pilot 2 | Google Patents BigQuery | 1,000 top-vernacular short list | none | +0.253 | $\approx 530$ | Negative $r$ at all four close-distance bins; $p = 0.001$ |
| Full run (main paper) | Google Patents BigQuery | All species mentioned in title/abstract | none | **+0.188** | 9,944 | Table 1 in main paper |

<sup>35</sup> Two observations drive the manuscript's conclusion that the result is not an artefact of corpus choice or sampling cap:

Two observations drive the manuscript’s conclusion that the result is not an artefact of corpus choice or sampling cap:

- **The three *r*-values are within 0.10 of each other** (range 0.188–0.281) and all have the same one-sided permutation *p* = 0.001 floor. Lifting the OPS top-25 cap (Pilot 1 → Pilot 2) moved *r* down by only 0.028 — the cap is not the source of the signal.
- **The correlogram shape is preserved across all three runs.** Every pilot shows negative *r* in the four close-distance bins ([0,3), [3,6), [6,9), [9,12)) and a sign-flip in the cross-kingdom bin ([12,∞)) — the same pattern reported in Table 1 of the main paper.

The full-tree run’s smaller global *r* = +0.188 (vs. +0.253 for the matched-shortlist BigQuery pilot) is expected: extending the species set from 530 well-studied taxa to 9,944 species across the full eukaryotic tree adds many cross-kingdom pairs (the [12, ∞) bin alone now holds 4.65 × 10^7^ of the 4.94 × 10^7^ total pairs, dominating the global Pearson correlation). This is precisely the dilution effect that motivates the correlogram decomposition reported in Table 1, in which each phylogenetic-distance bin’s signal is reported separately and remains highly significant at every close-distance bin.

### S2 — Supplementary Table S2: predictive query results (medicinal preparations, CPC *A61K*)

Results below were generated by running the Box 1 Cypher pattern against the live `treeoflife` graph with `$target = ’A61K’` and the maximum phylo-distance restricted to 2 (sister-species, same genus). Donor strength threshold: `$min_strength = 500`. For brevity, only the 15 strongest donors are shown; the full ranked list (700 donors at threshold 500, 3,691 at threshold 50) is reproducible from the query in Box 1. Candidates are species in the same genus as the donor that have at least one patent in the graph but no patent classified into *A61K*.

**Supplementary Table S2.**
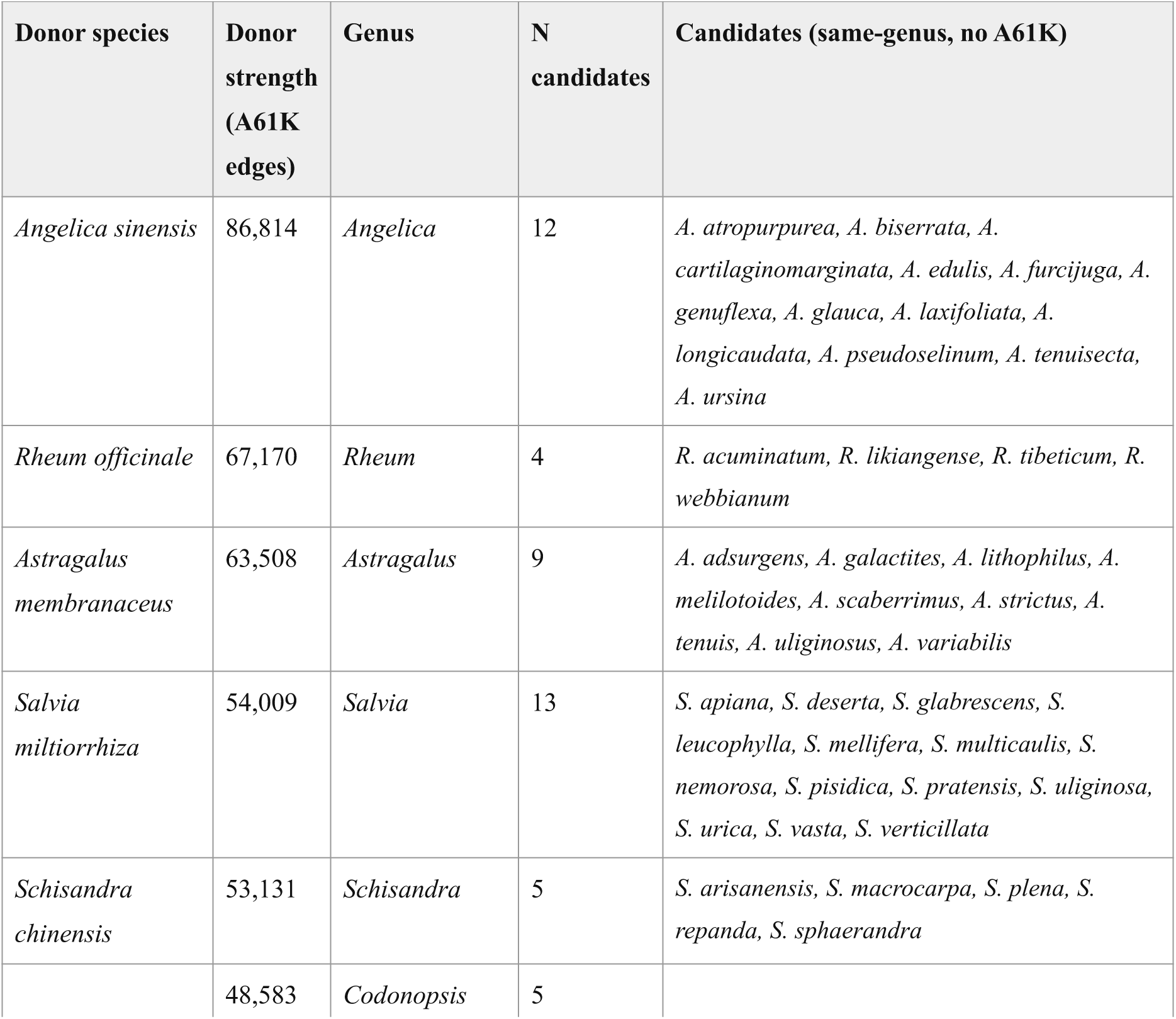
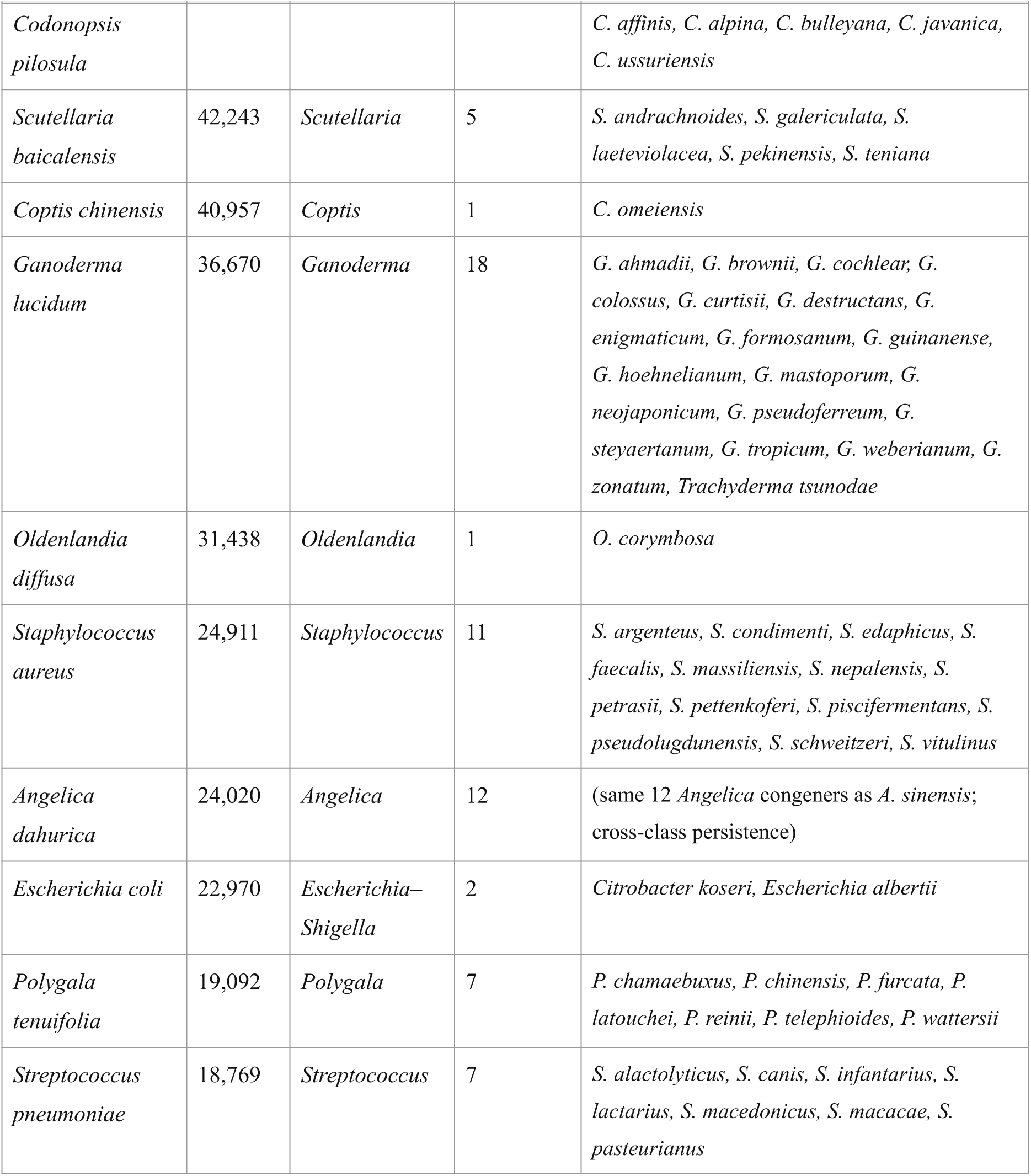
Top-15 A61K donors with sister-species candidates. Donor strength = number of `:HAS_PATENT`→*A61K*-classified edges from the donor species. Candidates listed in *italic*; all are at OTT phylogenetic distance 2 (same genus, no patent in the *A61K* subclass). Generated 2026-05-11 from the live graph.

### S3 — Supplementary Table S3: predictive query results (cosmetics, CPC *A61Q*)

As above, with `$target = ’A61Q’` and `$min_strength = 100`. The CPC *A61Q* subclass (specific use of cosmetics or similar toilet preparations) is structurally smaller than *A61K* (only 444 strong donors at threshold 50 vs. 3,691 for *A61K*), so the strength cutoff is lowered accordingly.

**Supplementary Table S3.** Top-10 A61Q donors with sister-species candidates. Donor strength = number of `:HAS_PATENT`→*A61Q*-classified edges from the donor species. Generated 2026-05-11.

| Donor species | Donor strength (A61Q edges) | Genus | N candidates | Candidates (same-genus, no A61Q) |
| --- | --- | --- | --- | --- |
| *Aloe vera* | 2,020 | *Aloe* | 9 | *A. africana*, *A. bella*, *A. humilis*, *A. maculata*, *A. nyeriensis*, *A. rauhii*, *A. schweinfurthii*, *A. spicata*, *A. succotrina* |
| *Staphylococcus aureus* | 1,464 | *Staphylococcus* | 22 | *S. argenteus*, *S. carnosus*, *S. chromogenes*, *S. condimenti*, *S. delphini*, *S. edaphicus*, *S. equorum*, *S. faecalis*, *S. kloosii*, *S. leei*, *S. massiliensis*, *S. muscae*, *S. nepalensis*, *S. pasteurii*, *S. petrasii*, *S. pettenkoferi*, *S. piscifermentans*, *S. pseudintermedius*, *S. pseudolugdunensis*, *S. schweitzeri*, *S. succinus*, *S. vitulinus* |
| *Bletilla striata* | 1,394 | *Bletilla* | 1 | *B. formosana* |
| *Lactobacillus plantarum* | 1,304 | *Lactobacillus* | 78 | 78 congeners; full list reproducible from the live query (top 5 shown): *L. acetotolerans*, *L. acidipiscis*, *L. agilis*, *L. algidus*, *L. alimentarius*... |
| *Ganoderma lucidum* | 1,181 | *Ganoderma* | 28 | 28 congeners; superset of the A61K candidates listed in Table S2. |
| *Glycyrrhiza glabra* | 1,148 | *Glycyrrhiza* | 4 | *G. aspera*, *G. echinata*, *G. glandulifera*, *G. korshinskyi* |
| *Scutellaria baicalensis* | 1,043 | *Scutellaria* | 18 | 18 congeners; superset of the A61K candidates listed in Table S2. |
| *Escherichia coli* | 937 | *Escherichia*–<br>*Shigella* | 5 | *E. albertii*, *E. fergusonii*, *Shigella boydii*, *S. flexneri*, *Citrobacter koseri* |
| *Gleditsia sinensis* | 886 | *Gleditsia* | 5 | *G. amorphoides*, *G. delavayi*, *G. fera*, *G. microphylla*, *G. triacanthos* |
| *Saccharomyces cerevisiae* | 811 | *Saccharomyces* | 9 | *S. bayanus*, *S. chevalieri*, *S. ellipsoideus*, *S. eubayanus*, *S. mikatae*, *S. minor*, *S. paradoxus*, *S. pastorianus*, *S. uvarum* |

### S4 — Cross-class consistency

Several donor genera appear in both Tables S2 and S3 (*Staphylococcus aureus*, *Ganoderma lucidum*, *Scutellaria baicalensis*, *Escherichia coli*), and their sister-species candidate sets overlap heavily — for example, all 11 *Staphylococcus* candidates flagged as A61K-unflagged are also flagged as A61Q-unflagged. The argument in the main paper (Example 2: "Cross-class consistency") is precisely this: when an identical sister-species is flagged by two independent application subclasses with two independent donor species in the same genus, the prior probability of the candidate being biotechnologically relevant rises sharply. The two-class intersection in this small slice already identifies ≈ 40 candidate species whose chemistry, physiology, or microbiome niche makes them simultaneous priors for both medicinal-preparation and cosmetics screening.

### S5 — Reproduction

To regenerate either table, run the Cypher pattern in **Box 1** of the main paper against the live graph with the parameters tabulated below.

| Parameter | Table S2 (A61K) | Table S3 (A61Q) |
| --- | --- | --- |
| \$target | 'A61K' | 'A61Q' |
| \$min_strength | 500 | 100 |
| Max phylo-distance | 2 (sister-species) | 2 (sister-species) |
| Result truncation | Top 15 donors | Top 10 donors |

*Graph commit:* 4,529,570 Taxon nodes, 759,182 Patent nodes, 35,139 CPCClass nodes — full-tree BigQuery overlay, build 2026-05-08. The graph state, the Cypher query, and the random seed (20260506) used by the Mantel pipeline together fully reproduce all numbers reported in the main paper and in this Supplementary Information.

### S6 — Branch-length (time-calibrated) analysis

To confirm that the phylogenetic signal is not an artefact of the unweighted OTT topology, we repeated the Mantel analysis using time-calibrated divergence times from the TimeTree of Life v5^10,11^ (Methods; main text Figure 4). Of the 9,944 patent-linked species, 5,387 (54.5%) matched a TToL5 tip by scientific name; these carry 72.4% of all patent volume in the graph, and 26 of the 30 most-patented species are present, so the matched subset is taxonomically thinner but retains most of the signal-bearing data. After removing 81 species whose patents carried only IPC codes, 5,306 species (14,074,165 pairs) entered the test. Phylogenetic distance was expressed as the divergence time (Myr) of each pair’s most-recent common ancestor on the exactly-ultrametric pruned tree (root age 3,772 Myr).

**Supplementary Table S4.** Branch-length Mantel correlogram by divergence-time bin (TToL5, n = 5,306 species, 999 permutations). Negative *r* indicates that pairs in the bin are more profile-similar than chance — the phylogenetic-signal direction. *p*_neg_ is the one-sided permutation *p*-value for negative *r*. *** = significant after Bonferroni at α = 0.05/7 = 0.007. Global Mantel *r* = +0.103, one-sided *p* = 0.001.

| Divergence-time bin (Myr) | Approx. taxonomic span | Pairs | $r$ | $p_{\text{neg}}$ | Verdict |
| --- | --- | --- | --- | --- | --- |
| [0, 10) | congeneric / recent | 6,017 | -0.027 | 0.001 | *** |
| [10, 50) | within-family | 39,575 | -0.018 | 0.001 | *** |
| [50, 100) | within-order | 173,630 | -0.022 | 0.001 | *** |
| [100, 250) | angiosperm / major-clade radiations | 3,542,204 | -0.105 | 0.001 | *** (peak) |
| [250, 500) | within-phylum | 735,334 | -0.034 | 0.001 | *** |
| [500, 1500) | cross-phylum | 3,586,536 | +0.068 | 1.000 | n.s. (anti-direction) |
| [1500, 4000) | cross-kingdom | 5,990,869 | +0.055 | 1.000 | n.s. (anti-direction) |

<sup>42</sup> The signal is Bonferroni-significant in every bin out to ~500 Myr of divergence and peaks at 100–250 Myr, the window of the angiosperm radiation and the diversification of the major animal and fungal groups that dominate the patent record. Beyond ~500 Myr the correlation flips positive and non-significant — the cross-kingdom dilution effect also seen in the topological $[12, \infty)$ bin of Table 1.

The signal is Bonferroni-significant in every bin out to ∼500 Myr of divergence and peaks at 100–250 Myr, the window of the angiosperm radiation and the diversification of the major animal and fungal groups that dominate the patent record. Beyond ∼500 Myr the correlation flips positive and non-significant — the cross-kingdom dilution effect also seen in the topological [12, ∞) bin of Table 1.

**Supplementary Table S5.** Metric-choice control: the result does not depend on topology vs time. All three Mantel tests use the same Jaccard distance on CPC subclasses; they differ only in the phylogenetic-distance metric and species set. The global correlation is positive and significant in every case.

| Distance metric | Tree | $n$ species | Global Mantel $r$ | one-sided $p$ |
| --- | --- | --- | --- | --- |
| Topological (edges through LCA) | OTT, full tree | 9,944 | +0.188 | 0.001 |
| Topological (edges through LCA) | TToL5, matched subset | 5,306 | +0.154 | 0.005 |
| Time-calibrated (divergence Myr) | TToL5, matched subset | 5,306 | +0.103 | 0.001 |

Recomputing the topological edge-count test on the identical 5,306-species TToL5 subtree (*r* = +0.154) brackets the time-calibrated estimate (*r* = +0.103) and the original full-tree topological value (*r* = +0.188): the conclusion is robust to whether phylogenetic distance is measured as topology or as evolutionary time, and to the change of underlying tree (OTT → TToL5). The smaller time-calibrated global *r* reflects the enormous Myr weights that deep cross-kingdom pairs receive on a 3,772-Myr-deep ultrametric tree, which dominate a single global Pearson correlation; the per-bin correlogram (Table S4) removes this leverage and is where the time-calibrated signal is sharpest.

### S7 — Web interface for interactive exploration

To let domain experts probe GRAFT without writing Cypher, we built a single-page browser served by a 200-line FastAPI back-end (`webapp/main.py`) wrapping four endpoints — `/api/taxon/search` (autocomplete on indexed `Taxon.name`), `/api/cpc` (CPC subclasses with patent and species counts), `/api/lineage` (root-ward path of any taxon), and `/api/tree` (the predictive subtree query in **Box 1** of the main paper, returning a forest of leaves plus up to six representative patent titles and Google Patents URLs per leaf, in one round-trip). The front-end (`webapp/static/{index.html, app.js, style.css}`) renders a D3 cluster dendrogram in vanilla JavaScript: species leaves are coloured green and labelled with their scientific binomial, English vernacular (where present in NCBI or GBIF), and patent count for the selected CPC subclass. Hovering a leaf reveals a tooltip with clickable patent titles linking to `patents.google.com` and a Wikipedia (en) link. Clicking an internal node re-roots the view at that node (drill down); a breadcrumb at the top shows the lineage of the current root, with each ancestor crumb clickable for drill-up.

**Worked example — Solanoideae × A61K.** Selecting the nightshade subfamily *Solanoideae* (OTT id 13867) and the medicinal-preparation subclass *A61K* returns 15 species ranked by patent count. The result tracks medicinal pharmacopoeia well outside the food-crop members of the family:

**Figure S1.**
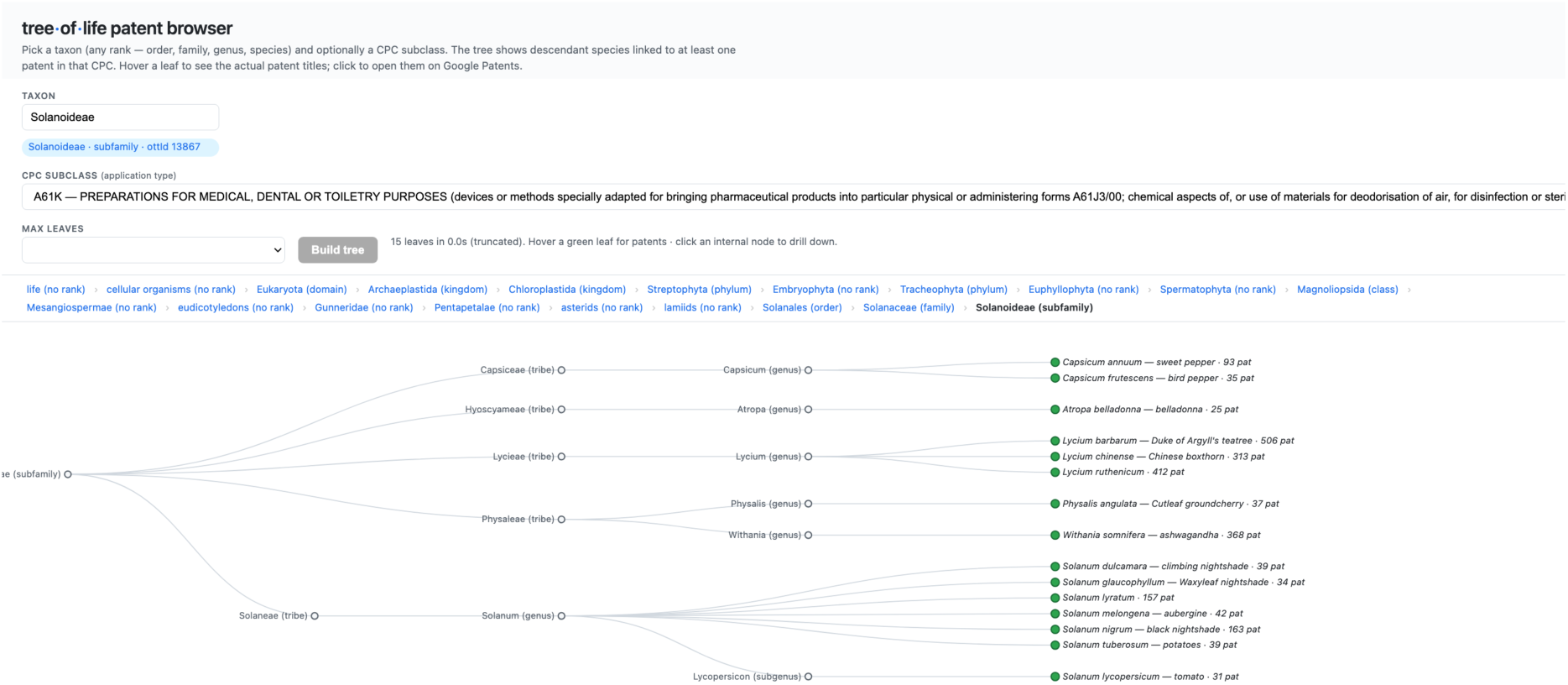
Web interface, Solanoideae × A61K view. Single-page browser at http://localhost:8000/?ott=13867&cpc=A61K&limit=15. The breadcrumb shows the lineage of the current root (*life › … › Solanaceae › Solanoideae*); each ancestor is clickable for drill-up. The dendrogram presents the 15 top-ranked nightshade species, each labelled with its scientific binomial, English vernacular (where available), and A61K-patent count. Mousing over any green leaf opens a tooltip with the patent titles linked to `patents.google.com` plus a Wikipedia link; clicking an internal node re-roots the view at that node. Output of the same Cypher pattern listed in Box 1 of the main paper.

**Supplementary Table S6.** Top-15 A61K-patented species in Solanoideae. (subfamily of Solanaceae). Direct output of the `/api/tree` endpoint with `ott=13867, cpc=A61K, limit=15`. Patent counts are distinct `:HAS_PATENT` edges; English vernacular pulled from GBIF / NCBI.

| Rank | Species | English name | A61K patents |
| --- | --- | --- | --- |
| 1 | *Lycium barbarum* | goji / Duke of Argyll's teatree | 506 |
| 2 | *Lycium ruthenicum* | (black goji) | 412 |
| 3 | *Withania somnifera* | ashwagandha | 368 |
| 4 | *Lycium chinense* | Chinese boxthorn | 313 |
| 5 | *Solanum nigrum* | black nightshade | 163 |
| 6 | *Solanum lyratum* | (no en. vernacular) | 157 |
| 7 | *Capsicum annuum* | sweet pepper | 93 |
| 8 | *Solanum melongena* | aubergine | 42 |
| 9 | *Solanum dulcamara* | climbing nightshade | 39 |
| 10 | *Solanum tuberosum* | potato | 39 |
| 11 | *Physalis angulata* | cutleaf groundcherry | 37 |
| 12 | *Capsicum frutescens* | bird pepper | 35 |
| 13 | *Solanum glaucophyllum* | waxyleaf nightshade | 34 |
| 14 | *Solanum lycopersicum* | tomato | 31 |
| 15 | *Atropa belladonna* | belladonna | 25 |

The ranking is dominated by traditional-medicine taxa rather than the food crops the family is famous for: the three *Lycium* species (goji berries) and *Withania somnifera* (Indian Ayurvedic ashwagandha) together carry 1,599 of the subfamily’s A61K patents — almost five times the combined total of *Solanum tuberosum* (potato), *S. lycopersicum* (tomato), *S. melongena* (aubergine), and *Capsicum annuum* (sweet pepper). The medical pharmacopoeia tags the historic medicinal genera even though the agricultural-economic weight sits with the food crops, an asymmetry the browser exposes at a glance.

Run the interface locally with `uvicorn webapp.main:app` --`reload` --`port 8000` after populating `.env`; the page is served at http://localhost:8000.

## References

1. Hinchliff, C. E., Smith, S. A., Allman, J. F. et al. Synthesis of phylogeny and taxonomy into a comprehensive tree of life. Proc. Natl Acad. Sci. USA 112, 12764–12769 (2015).

2. Schoch, C. L., Ciufo, S., Domrachev, M. et al. NCBI Taxonomy: a comprehensive update on curation, resources and tools. Database 2020, baaa062 (2020).

3. GBIF Secretariat. GBIF Backbone Taxonomy. Checklist dataset (2023). 10.15468/39omei.

4. Pagel, M. Inferring the historical patterns of biological evolution. Nature 401, 877–884 (1999).

5. Blomberg, S. P., Garland, T. Jr & Ives, A. R. Testing for phylogenetic signal in comparative data: behavioral traits are more labile. Evolution 57, 717–745 (2003).

6. Mantel, N. The detection of disease clustering and a generalized regression approach. Cancer Res. 27, 209–220 (1967).

7. Jaccard, P. The distribution of the flora in the alpine zone. New Phytol. 11, 37–50 (1912).

8. Legendre, P. & Fortin, M.-J. Spatial pattern and ecological analysis. Vegetatio 80, 107–138 (1989).

9. Lens.org. The Lens patent and scholarly database. https://www.lens.org (accessed May 2026).

10. Kumar, S., Suleski, M., Craig, J. M. et al. TimeTree 5: an expanded resource for species divergence times. Mol. Biol. Evol. 39, msac174 (2022).

11. Hedges, S. B., Marin, J., Suleski, M., Paymer, M. & Kumar, S. Tree of life reveals clock-like speciation and diversification. Mol. Biol. Evol. 32, 835–845 (2015).

